# The mitotic spindle protein CKAP2 potently increases formation and stability of microtubules

**DOI:** 10.1101/2021.07.20.453056

**Authors:** Thomas S. McAlear, Susanne Bechstedt

**Affiliations:** Department of Anatomy and Cell Biology, McGill University, Montreal, Canada

## Abstract

Cells increase microtubule dynamics to make large rearrangements to their microtubule cytoskeleton during cell division. Changes in microtubule dynamics are essential for the formation and function of the mitotic spindle, and misregulation can lead to aneuploidy and cancer. Using *in vitro* reconstitution assays we show that the mitotic spindle protein Cytoskeleton-Associated Protein 2 (CKAP2) has a strong effect on nucleation of microtubules by lowering the critical tubulin concentration 100-fold. CKAP2 increases the apparent rate constant *k_a_* of microtubule growth by 50-fold and increases microtubule growth rates. In addition, CKAP2 strongly suppresses catastrophes. Our results identify CKAP2 as the most potent microtubule growth factor to date. These finding help explain CKAP2s role as an important spindle protein, proliferation marker, and oncogene.

## Introduction

During mitosis, cells build mitotic spindles to faithfully segregate their chromosomes. Assembling mitotic spindles requires a complete rearrangement of the microtubule cytoskeleton, which is driven by the concerted action of microtubule-associated proteins (MAPs) and motor proteins (Kapoor, 2017). Microtubule turnover increases significantly in mitosis resulting from increased microtubule nucleation (Piehl et al., 2004) in combination with shorter microtubule lifetimes (Saxton et al., 1984).

How exactly microtubule nucleation and growth are mediated in mitotic spindles and if the major assembly factors have been identified remains an open question. XMAP215/chTOG and TPX2 have been implicated as two major factors in nucleation (Roostalu et al., 2015). In addition, XMAP215/chTOG family members are the only protein shown to speed up microtubule growth by more than a factor of 3 in *in vitro* assays (Brouhard et al., 2008).

Microtubules nucleate primarily from centrosomes as well as several centrosome independent microtubule nucleation pathways (Petry and Vale, 2015). Recently, it has been discovered that new microtubules can directly nucleate from pre-existing microtubules through augmin-mediated branched microtubule nucleation (Petry et al., 2011; Uehara et al., 2009). Most nucleation pathways include γ-tubulin ring complexes (γ-TURCs), multi-protein assemblies that are thought to promote nucleation and may also function to cap the microtubule minus ends. Notably, the γ-TURC only weakly increases microtubule nucleation and is thought to require activation of the complex itself or the help of MAPs like the microtubule polymerase XMAP215/chTOG or the nucleation factor TPX2 (Consolati et al., 2020; Kollman et al., 2010; Moritz et al., 1995; Thawani et al., 2018).

Given the robustness of spindle assembly as well as the presence of microtubules in cells lacking chTOG (Gergely et al., 2003), TPX2 (Aguirre-Portolés et al., 2012), and γ-tubulin (Strome et al., 2001), it seems likely that additional factors can promote microtubule formation in spindles.

The Cytoskeleton-Associated Protein 2 (CKAP2) is an intrinsically disordered protein (Figure 1A) that localizes to mitotic spindles during cell division (Seki and Fang, 2007). CKAP2 expression and phosphorylation states are tightly regulated throughout the cell cycle. During G1 interphase, CKAP2 expression is at the detection level and increases at the onset of mitosis (Seki and Fang, 2007). CKAP2 is regulated by sharp phosphorylation events between different mitotic stages (Hong et al., 2009). The Anaphase-promoting complex (APC/C) marks CKAP2 for degradation at the end of mitosis to eliminate all CKAP2 from the newly formed daughter cells (Hong et al., 2007).

**Figure 1.**
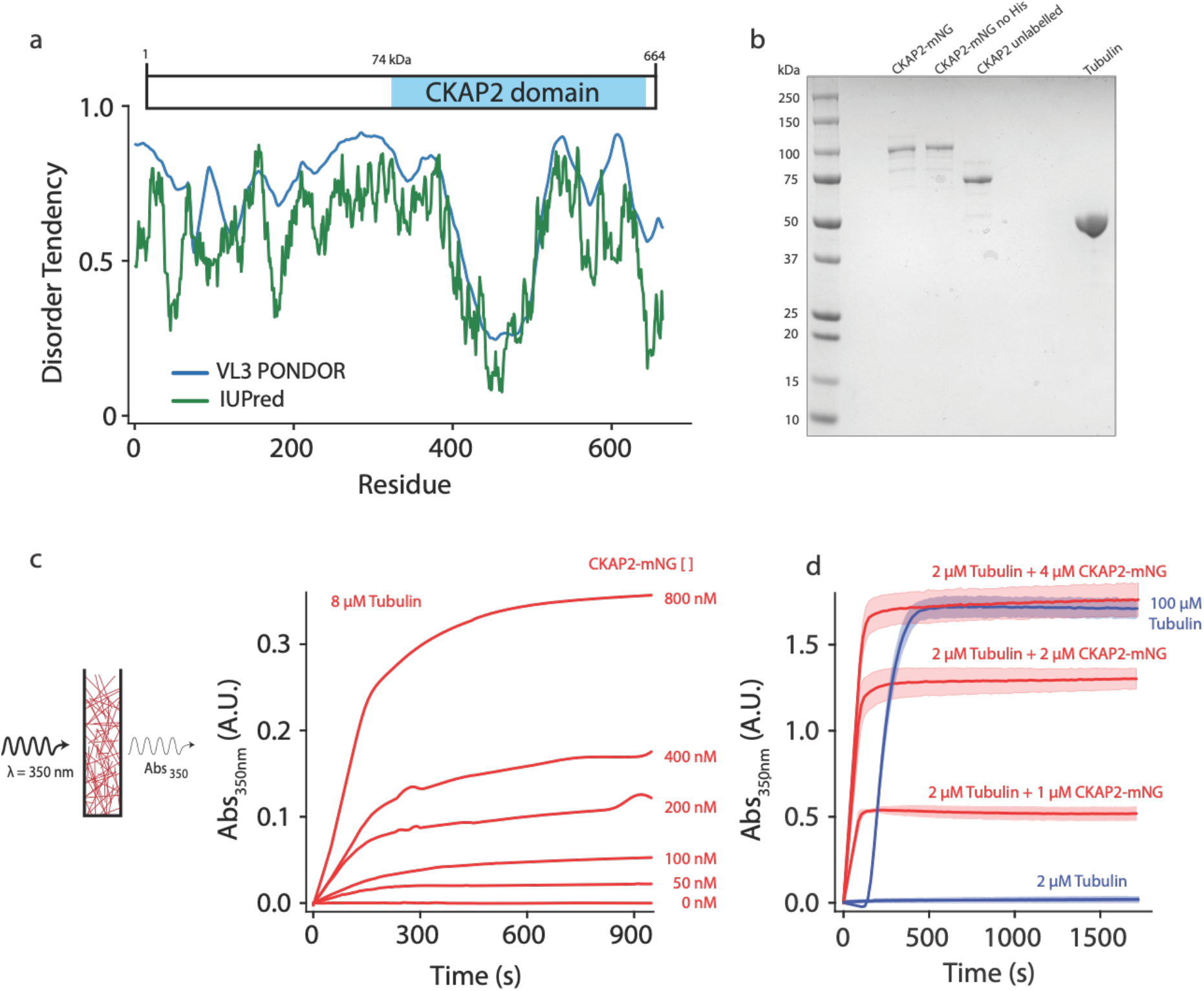
CKAP2 is an intrinsically disordered protein that increases microtubule formation. **(a)** Schematic of mmCKAP2 protein domains and disorder prediction (Dosztányi et al., 2005). **(b)** Coomassie Blue-stained SDS-PAGE gel of 1 μg of purified recombinant CKAP2 constructs and 5 μg of purified tubulin. **(c)** Light scattering assay (turbidity) schematic and data following microtubule formation as apparent absorbance (Abs) over time for 8 μM tubulin with increasing concentrations of CKAP2-mNeonGreen (CKAP2-mNG) (*n* = 1). **(d)** Turbidity data for tubulin alone (blue) and addition of CKAP2-mNG (*n* = 3, mean ± SD).

Knock-down of CKAP2 in cells interferes with proper spindle assembly and often results in multipolar spindles and misaligned chromosomes (Case et al., 2013). Overexpression of CKAP2 promotes cancer formation (Guo et al., 2017; Yu et al., 2015) and correlates with severity of the disease (Hayashi et al., 2014). Consequently, CKAP2 is a marker for the diagnosis and prognosis of several types of cancer (Hayashi et al., 2014). CKAP2 has been previously described as a potential microtubule stabilizer in cells (Jin et al., 2004; Tsuchihara et al., 2005), though it is not known whether CKAP2 directly binds to microtubules or how it affects microtubule dynamics. We were intrigued by the cell biology of CKAP2, the severe spindle phenotypes, the clinical relevance, and the lack of any molecular understanding of this protein. Therefore, we wanted to investigate how CKAP2 impacts microtubule dynamics.

## Results

To test for the ability of CKAP2 to influence microtubule polymer formation, we used recombinantly expressed full-length protein (Figure 1B) in a bulk light scattering assay (Figure 1C). At physiological tubulin levels (8 μM), increasing amounts of CKAP2 cause dose-dependent increase in light scattering and apparent absorbance (turbidity). At higher concentrations of CKAP2 turbidity continued to increase (Figure 1D). Without CKAP2, 50-fold more tubulin was necessary to reach the same turbidity observed for 2 μM tubulin in the presence of 4 μM CKAP2 (Figure 1D). Additionally, CKAP2 increased turbidity at 4 °C (Figure S1C), a temperature where tubulin does not form polymers and preformed microtubules depolymerize quickly in the absence of any stabilizers. Neither the mNeonGreen (mNG) nor the 6-His affinity tag had any impact on the ability of CKAP2 to aid microtubule formation (Figures S1A and S1B).

The bulk turbidity assay cannot distinguish between effects of CKAP2 on microtubule nucleation, growth, or stabilization. To test how CKAP2 enhances tubulin polymer formation, we reconstituted microtubule assembly *in vitro* using total internal fluorescence (TIRF) microscopy (Gell et al., 2010). In this assay we observe individual microtubules growing from GMPCPP stabilized microtubule ‘seeds’ (Figure 2A). At physiological tubulin concentrations and in the presence of CKAP2, observation of individual growth events were only possible at very low CKAP2 concentrations (≤50 nM) due to high levels of spontaneous nucleation beyond that point. The microtubule growth rate increased linearly with CKAP2 at these concentrations (Figures 2B and S2A).

**Figure 2.**
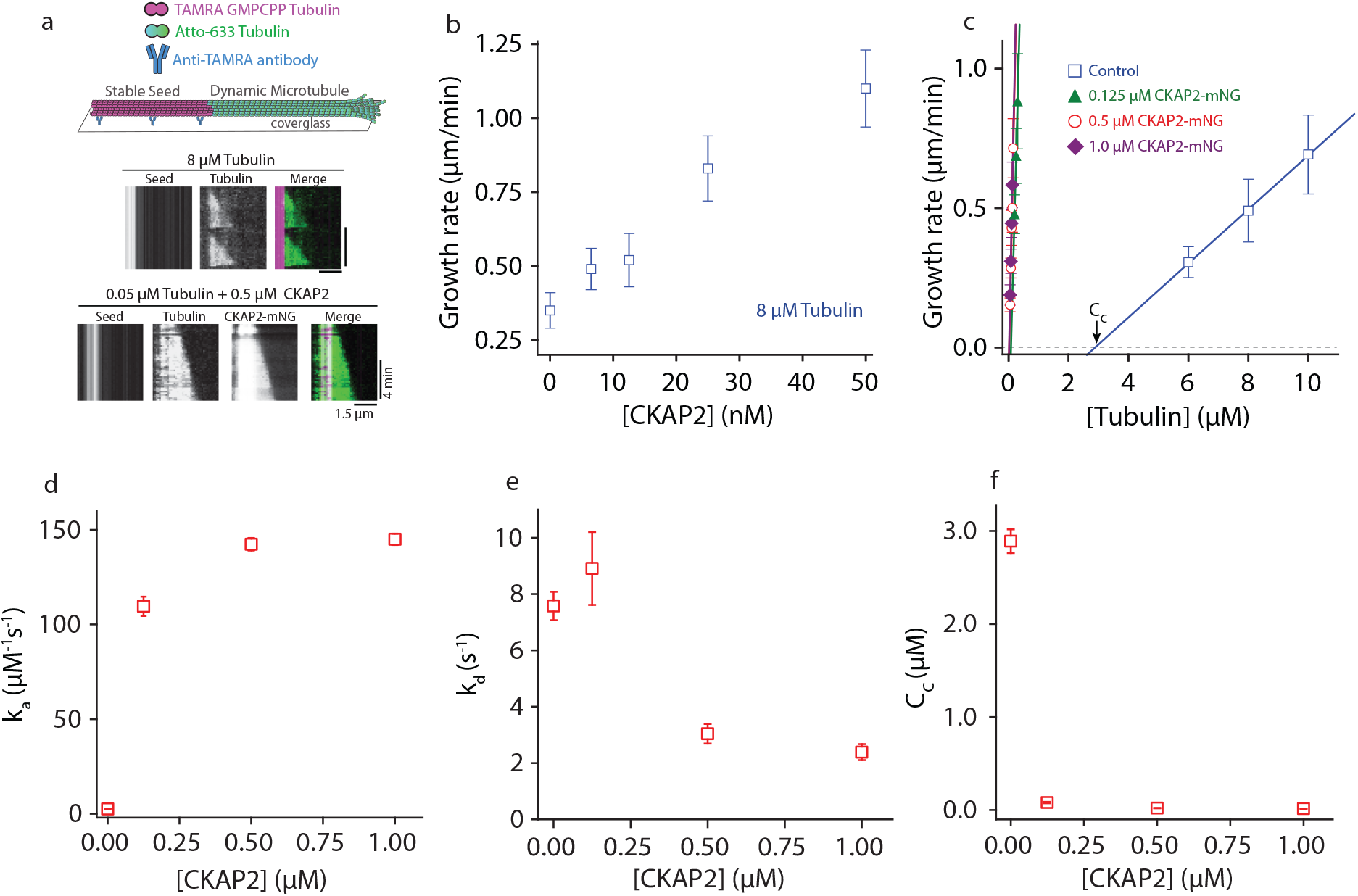
CKAP2 lowers the critical concentration of microtubule growth and speeds up microtubule assembly rates. **(a)** Dynamic microtubule growth assay schematic (upper panel). Microtubule dynamics for tubulin control and in the presence of CKAP2-mNG are analyzed from space-time plots (kymographs, lower panel). **(b)** Plot of microtubule growth rates as a function of CKAP2-mNG concentration with 8 μM tubulin. Plotted as mean ± SD (*n* = 67, 24, 70, 67, 46 from 1-2 replicates) **(c)** Plot of microtubule growth rates as a function of tubulin concentration for control (no CKAP2) and in the presence of CKAP2-mNG. Plotted as mean ± SD (blue; *n* = 272, 601, 347 from 3 replicates), 0.125 μM CKAP2-mNG (green; *n* = 146, 85, 91 from 2 replicates), 0.5 μM CKAP2-mNG (red; *n* = 48, 92, 91, 85, 55 from 2 replicates) and 1 μM CKAP2-mNG (purple; *n* = 68, 134, 143, 115 from 2 replicates). **(d)** Plot of the apparent on-rate constant (*k_a_*) as a function of CKAP2-mNG concentration determined from linear regression fit of growth rates versus tubulin concentration. Error bars represent SE of fit. **(e)** Plot of the apparent off-rate constant (*k_d_*) as a function of CKAP2-mNG concentration determined from linear regression fit of growth rates versus tubulin concentration. Error bars represent SE of fit. **(f)** Plot of the apparent critical concentration (*C_c_*) as a function of CKAP2-mNG concentration determined from linear regression fit of growth rates versus tubulin concentration. Error bars represent SE of fit.

To measure microtubule growth across a wide range of CKAP2 concentrations and determine apparent tubulin on-and off rates (k_a_ and k_d_) we reduced the tubulin concentration about 100-fold (to 50-300 nM) to mitigate spontaneous nucleation and observe individual microtubule growth events. The resulting growth curves (Figures 2C and S2B) illustrate microtubule elongation at substantially lower tubulin concentrations than previously observed. Microtubules polymerized with an apparent assembly rate constant (*k_a_*) of 2.6 ± 0.06 dimers·μM^−1^·sec^−1^ in controls and 142 ± 3.3 dimers·μM^−1^·sec^−1^ in the presence of 500 nM CKAP2, representing a 54-fold increase in *k_a_* (Figure 2D). By comparison, members of the XMAP215/chTOG family of microtubule polymerases achieve about 5-fold maximal increase in *k_a_* (Brouhard et al., 2008). We observed a 42-fold increase in *k_a_* for 125 nM CKAP2. Increasing the CKAP2 concentration beyond 500 nM did not further increase growth rates. We observe a smaller effect (∼ 3-fold) on the tubulin off-rate k_d_ (Figure 2E).

From the growth curves in Figure 2C and the spontaneous nucleation we observe at near-physiological concentrations of CKAP2 and tubulin, it is apparent that CKAP2 is able to shift the critical concentration (Cc) for microtubule elongation into the low nanomolar range (from 2.89 ± 0.12 μM to 0.02 ± 0.002 μM with 500 nM CKAP2) (Figure 2F).

To characterize the nucleation behaviour of microtubules we turned to an assay for templated nucleation from seeds (Wieczorek et al., 2015) (Figure 3A). We measured the probability of a seed to nucleate a microtubule over time. At physiological tubulin levels microtubules nucleated faster in the presence of as ≥25 nM CKAP2 (Figure 3B). When we explored a wider range of CKAP2 and tubulin concentrations, we found a very strong promotion of templated nucleation by CKAP2 even at 100-fold lower tubulin concentrations compared to controls (Figure 3C, S3A and S3E). The tubulin concentration for the half-maximal probability to nucleate within one minute is reduced from 6.85 ± 0.48 μM in controls to 0.05 ± 0.01 μM for 0.5 μM CKAP2-mNG. CKAP2 facilitates templated nucleation from seeds at tubulin concentrations as low as 50 nM. Further, we quantified the spontaneous nucleation, i.e. in the absence of templates, in our microscopy assay by counting the number of microtubules formed after two minutes (Figure 3D) (Roostalu et al., 2015). We found that CKAP2 dramatically increases spontaneous nucleation with a shift in critical tubulin concentration from 25.4 ± 1.7 μM to 0.25 ± 0.02 μM (Figure 3E and S3B). CKAP2 therefore promotes both templated as well as spontaneous nucleation. In the presence of CKAP2 the critical concentration of spontaneous nucleation, as well as nucleation from templates, are lowered up to 100-fold.

**Figure 3.**
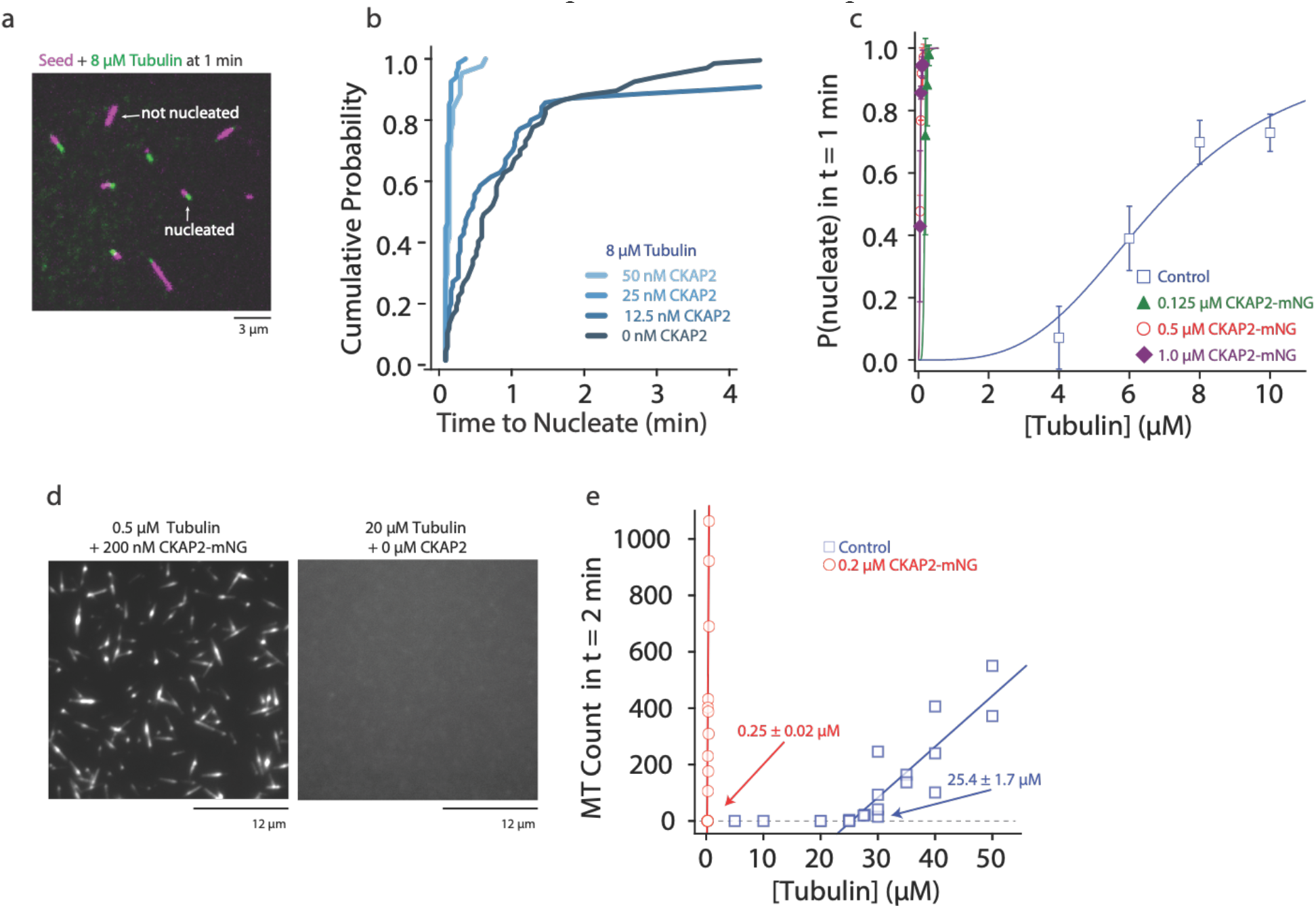
CKAP2 increases templated and spontaneous microtubule nucleation, and suppresses catastrophe. **(a)** Representative field of view of GMPCPP-stabilized microtubule seeds nucleating microtubules from 8 μM tubulin. **(b)** Plot of the cumulative probability distributions for time to nucleate as a function of CKAP2-mNG concentration (*n* = 67, 70, 67, 46 from 2 replicates) **(c)** Plot of probability of GMPCPP seed to be nucleated within 1 minute for control (blue; n = 102, 271, 115, 184 from ≥2 replicates), 0.125 μM CKAP2-mNG (green; *n* = 164, 213, 166 from 2 replicates), 0.50 μM CKAP2-mNG (red; *n* = 179, 216, 145, 126, 145 from 2 replicates) and 1 μM CKAP2-mNG (purple; *n* = 112, 186, 182, 148 from 2 replicates). Plotted as mean ± SD. Data was fit to a hill function forced to start at y=0 and end at y=1 (y = START + (END - START) * x^n / (k^n + x^n)). Tubulin concentrations for half-maximal microtubule growth, C, and steepness of fit, s, for control (C = 6.85 ± 0.48 μM, s = 3.37 ± 0.87) 0.125 μM CKAP2-mNG (C = 0.18 ± 0.01 μM, s = 6.96 ± 1.39) 0.5 μM CKAP2-mNG (C = 0.05 ± 0.01 μM, s = 3.10 ± 0.20) and 1 μM CKAP2-mNG (C = 0.05± 0.01 μM, s = 3.79 ± 0.85) **(d)** Representative field of view of microtubules spontaneously nucleated in the absence of templates. **(e)** Quantification of spontaneous nucleation and linear fit showing a critical tubulin concentration for microtubule nucleation of 25.4 ± 1.7 μM for tubulin alone (blue; *n* = 20) and 0.25 ± 0.02 μM in the presence of 0.2 μM CKAP2 and 0.25 ± 0.02 μM (red; *n* = 14).

A hallmark of microtubule dynamic instability is their capacity to spontaneously switch from growth to shrinkage, a behavior termed catastrophe (Mitchison and Kirschner, 1984). When microtubules were grown in the presence of CKAP2, catastrophe levels were severely reduced (Figures 4A). We quantified this effect and determined that CKAP2 lowers catastrophe frequencies in a concentration dependent manner, reducing them to near zero at 50 nM CKAP2 (Figure 4B). In the presence of higher concentrations of CKAP2, catastrophe frequency was at least an order of magnitude lower than in controls (Figure 4C, S4A, S4B and S4C). The only other family of MAPs with a comparable effect on microtubule catastrophes are the cytoplasmic linker associated proteins (CLASPs) (Aher et al., 2018; Lawrence et al., 2018; Majumdar et al., 2018).

**Figure 4.**
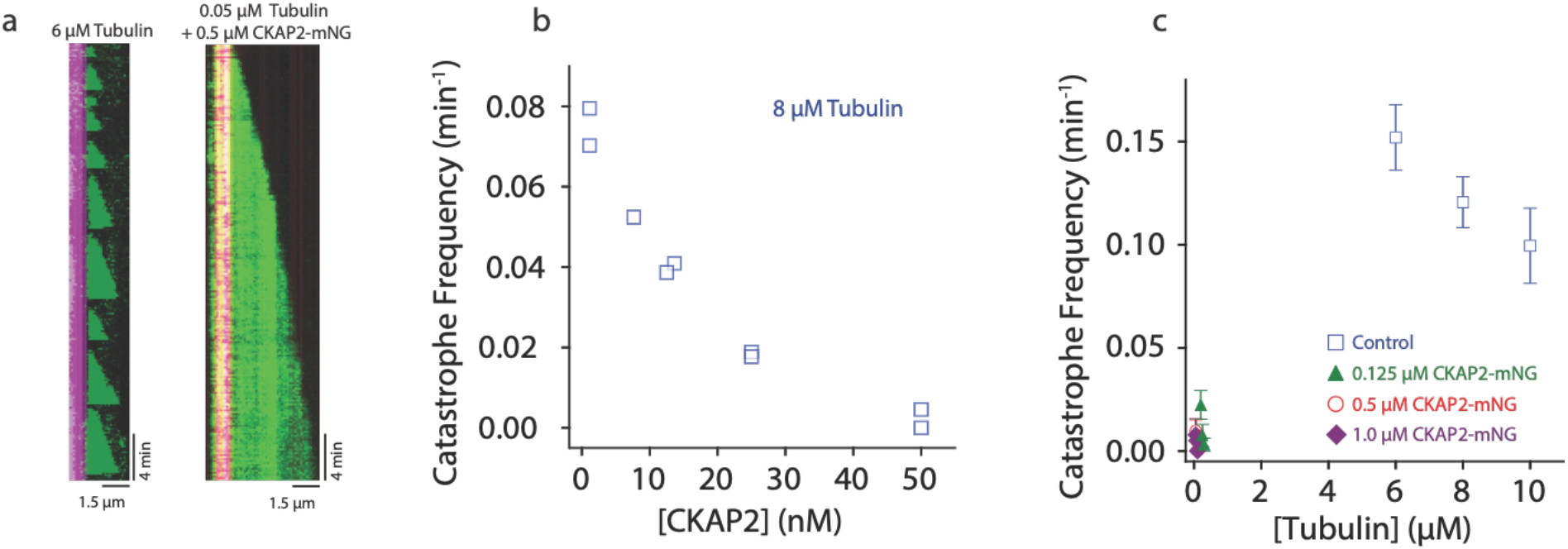
CKAP2 suppresses catastrophe. **(a)** Kymographs of representative microtubule growth in the absence and presence of CKAP2-mNG. **(b)** Plot showing the microtubule catastrophe frequencies as a function of CKAP2-mNG concentration. Each point represents the catastrophe frequency for an individual channel. (*n* = 19-39 microtubule growth phases for 0 nM to 50 nM) **(c)** Plot showing the microtubule catastrophe frequencies as a function of tubulin concentration for control (blue; *n* = 217, 676, 502 from ≥2 replicates), 0.125 μM CKAP2-mNG (green; *n* = 133, 222, 166 from 2 replicates), 0.50 μM CKAP2-mNG (red; *n* = 234, 232, 95, 135 from 2 replicates) and 1 μM CKAP2-mNG (purple; (*n* = 75, 171, 182, 148 from 2 replicates). Plotted as mean ± SD

A prediction for a potential cytoskeletal growth factor is the interaction with both the growing filament end as well as free monomers, as it has been shown for members of the chTOG/XMAP215 family for microtubules (Ayaz et al., 2014) and formins for actin (Kovar et al., 2006). We tested this hypothesis directly by assaying if CKAP2 is able to recruit soluble tubulin to microtubules. Indeed, lattice bound CKAP2 recruits free Atto-663 tubulin to GMPCPP microtubule seeds (Figure 5A and B). CKAP2 therefore possesses a distinct microtubule lattice as well as likely one or multiple tubulin binding sites. Recruitment of tubulin by CKAP2 could therefore play a role in promoting microtubule nucleation and growth.

**Figure 5.**
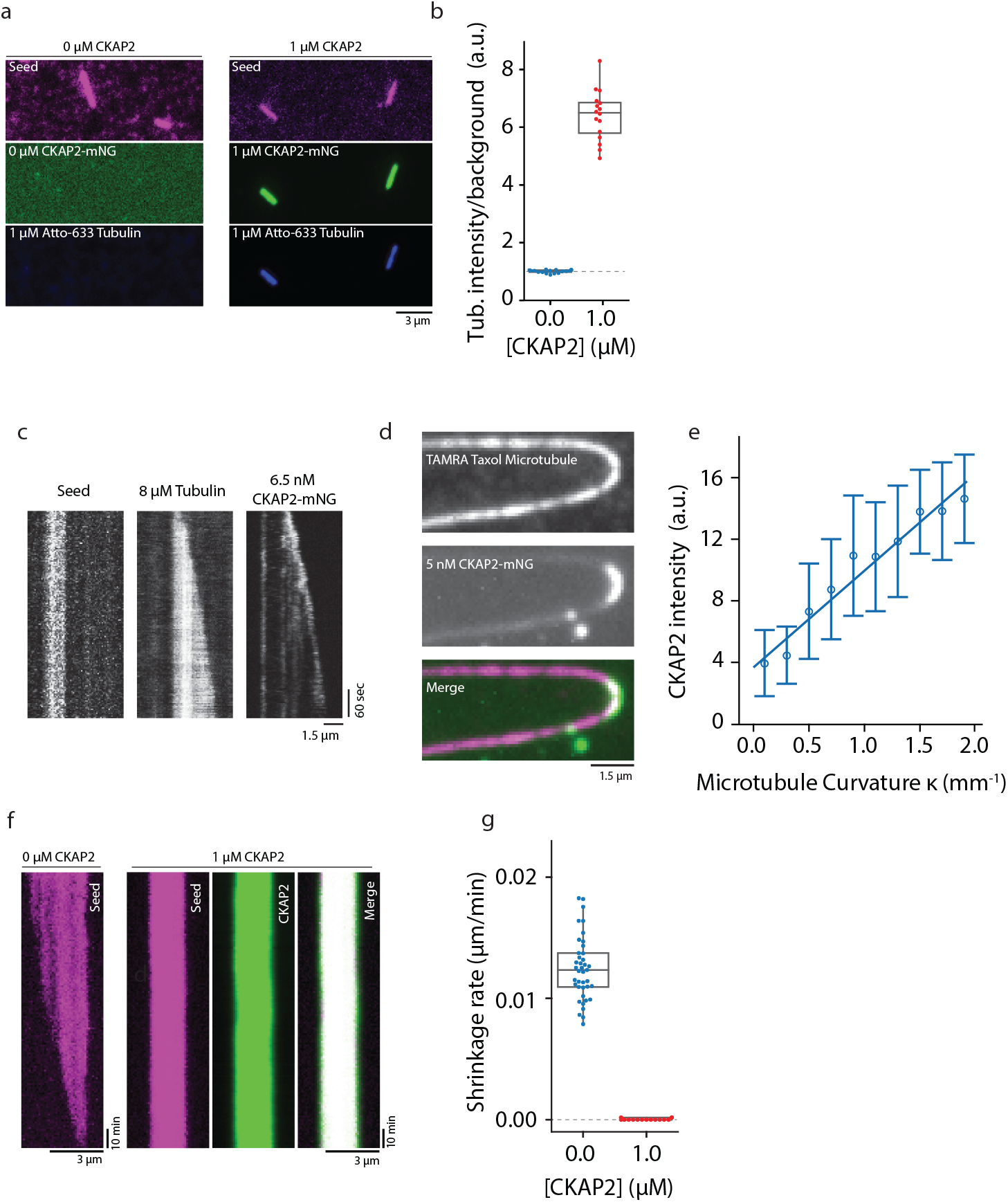
CKAP2 recruits tubulin, recognizes lattice curvature, but does not catalyze microtubule depolymerization. **(a)** Field of view showing 1 μM CKAP2-mNG recruiting Atto-633 tubulin to GMPCPP seeds. **(b)** Box plot of tubulin intensity on the GMPCPP lattice over background for control and 1 μM CKAP2-mNG (blue; *n* = 17 from 1 replicate) and 1 μM CKAP2-mNG (red; *n* =16 from 1 replicate). **(c)** Kymograph representative of end-tracking events of 6.5 nM CKAP2-mNG on a growing microtubule end. **(d)** Field of view of 5 nM CKAP2-mNG preferentially binding to curved regions of TAMRA-labeled paclitaxel-stabilized microtubules. **(e)** Plot of CKAP2-mNG intensity versus curvature (κ) measured using Kappa (Mary and Brouhard, 2019) (*n* = 73 from 2 replicates. Plotted as binned mean ± SD) **(f)** GMPCPP seed depolymerization kymographs representative of *n* = 41 (control) and *n* = 16 (1 μM CKAP2-mNG). **(g)** Box plot displaying the shrinkage rate of GMPCPP microtubule seeds for control (blue; n = 41 from 1 replicate) and in the presence of 1 μM CKAP2-mNG (red; *n* =16 from 1 replicate).

Many of the proteins promoting microtubule nucleation and growth possess high affinity binding sites at the plus end of dynamic microtubules (e.g. XMAP215/chTOG, DCX, TPX2). Here, XMAP215/chTOG is thought to help the addition of tubulin dimers to the growing end (Howard and Hyman, 2007). To determine if CKAP2 interacts with the growing microtubule end, we performed our TIRF microscopy assay at sub-saturating CKAP2 concentrations. Indeed, at low protein concentrations (6.5 nM), CKAP2 tracks microtubule ends (Figure 5C). At higher concentrations end tracking is obscured by lattice binding reminiscent of end tracking behavior observed for the microtubule nucleators DCX (Bechstedt and Brouhard, 2012) and TPX2 (Roostalu et al., 2015). Similar to TPX2 (Roostalu et al., 2015) and DCX (Bechstedt and Brouhard, 2012), CKAP2 recognizes microtubule curvature (Figures 5D and E), which is thought to correlate with the high affinity binding site of these proteins at the microtubule end (Bechstedt et al., 2014; Roostalu et al., 2015). In contrast, members of the XMAP215/chTOG family display robust autonomous end tracking at all physiologically relevant concentrations and do not recognize lattice curvature (Brouhard et al., 2008; Roostalu et al., 2015).

We next tested if CKAP2 can promote the removal of tubulin dimers from the lattice in the absence of tubulin. Catalyzing this reverse reaction of microtubule growth is a hallmark of *bona fide* microtubule polymerases of the XMAP215/chTOG family (Brouhard et al., 2008). We did not observe any depolymerization over ∼10 hrs (Figures 5F,G and S5A) when we incubated GMPCPP-stabilized microtubules with CKAP2 at concentrations as high as 1 μM, while controls without CKAP2 displayed an expected depolymerization rate of about 0.013 ± 0.003 μm/min (Figures 5F and G). CKAP2 does not catalyze the reverse reaction. On the contrary, CKAP2 strongly reduces the tubulin off-rate.

## Discussion

Our results show that CKAP2 has a strong effect on microtubule growth, nucleation, and stabilization in our *in vitro* system.

Cellular concentrations of CKAP2 in *Xenopus* eggs have been measured at about 70 nM (Wühr et al., 2014). In the same system the tubulin concentration was determined to be ∼ 7.5 μM. At physiological CKAP2 and tubulin concentrations (assuming 70 nM and 8 μM) CKAP2 has a strong effect on the microtubule growth rate (∼5-fold) in our assays. Further, templated nucleation is almost instantaneous and spontaneous nucleation is detectable. Catastrophe frequency is approaching zero at this concentration regime. Therefore, in cells, CKAP2 could increase microtubule nucleation, growth rates, and suppress catastrophe. Interestingly, at cellular CKAP2 concentrations, the effects on microtubule dynamics in our *in vitro* assays are not fully saturated. Hence, cells may be susceptible to increased levels of CKAP2 as they have been observed in several types of cancer.

Notably, we purify CKAP2 from *E.coli* in an unphosphorylated state. In cells, sharp phosphorylation events are observed for CKAP2 between different mitotic stages which could modify and shift the effect of CKAP2 on different microtubule dynamics parameters (Hong et al., 2008).

At full biochemical capacity of CKAP2, we observed a 100-fold increase in spontaneous and templated nucleation as well as a 54-fold increase in apparent tubulin on rate k_a_, and a complete suppression of microtubule catastrophe. Combined, these effects result in CKAP2 being the most potent microtubule assembly factor characterized to date.

Mechanistically, CKAP2 combines characteristics of microtubule nucleators (TPX2, DCX) as well as polymerases (XMAP215/chTOG) and anti-catastrophe factors (CLASPs). Like TPX2 and DCX, CKAP2 aids spontaneous nucleation. All three proteins display end tracking behaviour at low concentrations as well as lattice curvature recognition. These characteristics seem to be a hallmark of microtubule nucleation factors and might point towards the structural nature of the nascent nucleus. Similar to TPX2, CKAP2 dampens tubulin off-rates and microtubule dynamicity (Reid et al., 2016; Roostalu et al., 2015). A potential mechanism for nucleation by CKAP2 is the stabilization of the nascent microtubule nucleus (Roostalu and Surrey, 2017). Similar to the CLASP anti-catastrophe factors, CKAP2 lowers the catastrophe frequency to virtually zero (Aher et al., 2018; Lawrence et al., 2018; Majumdar et al., 2018), allowing for continued microtubule growth even at nanomolar concentrations of tubulin.

Other MAPs that stabilize the microtubule lattice like DCX (Moores et al., 2006), TPX2 (Reid et al., 2016), and CLASPs (Lawrence et al., 2018; Yu et al., 2016), as well as stabilizing drugs like paclitaxel (Zanic et al., 2013) or GTP analogs like GMPCPP (Hyman et al., 1992), do not severely impact the microtubule growth curve. Therefore, lattice stabilization alone cannot account for the severe impact of CKAP2 on the apparent tubulin on rate k_a_.

Unlike TPX2, DCX, and CLASPs, CKAP2 also acts, in part, like a polymerase. Similar to known polymerases of the XMAP215/chTOG family, CKAP2 recruits tubulin and increases the association rate constant (k_a_) specifically at the microtubule plus end (Brouhard et al., 2008; Gard and Kirschner, 1987). In contrast to XMAP215/chTOG family members, CKAP2 is able to push microtubule growth to the physical limit as defined by diffusion of the tubulin dimer to the growing microtubule end (Odde, 1997; Zanic et al., 2013). Unlike a polymerase, CKAP2 does not catalyze the reverse reaction of tubulin polymerization in the absence of tubulin substrates. CKAP2 either does not operate as a *bona fide* polymerase, or the depolymerase activity is masked by the lattice stabilization activity of CKAP2.

In many ways microtubule growth in the presence of CKAP2 resembles that of microtubules grown in the presence of the slowly hydrolysable GTP analog, GMPCPP. With GMPCPP, microtubules nucleate at nanomolar concentrations of tubulin and are relatively stable (Desai and Mitchison, 1997; Hyman et al., 1992). However, GMPCPP does not increase apparent on rate k_a_, like CKAP2.

How does CKAP2 combine multiple functionalities? The protein is about one third of the size of XMAP215/chTOG, about half the size of CLASPs, and is lacking any known microtubule or tubulin binding domains. We believe that the highly disordered nature of CKAP2 (Figure 1A) contributes to its ability to dynamically interact with the microtubule lattice as well as tubulin and impact microtubule assembly to an extreme extent. The advantage of disordered over folded domains is the absence of a structured, tight binding conformation to a substrate or intermediate state. In case of a microtubule growth factor, lack of a fixed structure would allow for fast molecular recognition of the tubulin dimer as well as for rapid unbinding, i.e. letting go of the dimer when incorporated (Uversky and Dunker, 2013). Structural as well as structure-function studies are needed to determine the interaction domains with microtubules and tubulin and provide us with further mechanistic insight.

Even small changes in microtubule assembly rates are correlated with increased levels of chromosomal instability (CIN) in cells (Ertych et al., 2014). CIN can provide the high adaptation capability necessary for tumor initiation and progression. Correspondingly, it has been shown that reducing microtubule growth rates in cancer cells by chemical or genetic manipulation suppresses CIN (Ertych et al., 2014). Misregulation of a potent growth factor like CKAP2 as observed in many cancers can therefore be directly responsible for CIN, and subsequently for aneuploidy, and malignancies.

Therefore, cells need to tightly regulate the capacity of a potent microtubule growth factor like CKAP2. Known regulatory measures include rapid degradation by APC (Seki and Fang, 2007) as well as distinct phosphorylation and dephosphorylation events in between different stages of the cell cycle (Hong et al., 2008, 2009). Phosphorylation could alter the effects of CKAP2 on microtubule dynamics and tune CKAP2 functionality between nucleation, stabilization, and growth.

Together, our findings identify the mitotic spindle protein CKAP2 as the most potent microtubule growth factor to date. The protein displays unprecedented impact on microtubule growth, nucleation, and catastrophe. Our results suggest an explanation for the observed CKAP2 knock-down spindle phenotypes and the oncogenic potential through the misregulation of microtubule dynamics.

## Materials and Methods

### Protein Expression and Purification

The coding sequence for full-length mouse CKAP2 protein (Uniprot accession # Q3V1H1) was PCR amplified from cDNA from wild-type mouse testes (a gift from Alana Watt’s Lab) using PfuX7 (Norholm, 2010) polymerase and inserted into a modified pHAT vector containing an N-terminal 6xHis-tag with and without a carboxy-terminal mNeonGreen (Allele Biotech) followed by a Strep-tag II (Bechstedt and Brouhard, 2012). Full length CKAP2-without a 6xHis-tag still containing a carboxy-terminal mNeonGreen followed by a Strep-tag II was synthesized into a pET21 vector (Twist Bioscience). For recombinant protein expression, BL21(DE3) *E. coli* containing protein expression vectors were grown to OD 0.6 at 37 °C, and expression was induced using 0.5 mM IPTG at 18 °C for 16 h. Bacterial pellets were harvested by centrifugation and resuspended in Buffer A (50 mM Na2HPO4, 300 mM NaCl, 4 mM Imidazole pH 7.8). Cells were lysed using a French press (EmulsiFlex-C5, Avestin). Constructs containing both His and Strep tags were purified using gravity flow columns containing His60 Ni-NTA resin (Clontech) followed by Streptactin affinity chromatography (IBA Lifesciences, Germany). Purified CKAP2 was eluted with BRB80 (80 mM PIPES-KOH pH 6.85, 1 mM EGTA, 1 mM MgCl_2_) containing 2.5 mM desthiobiotin and 10% glycerol. For the CKAP2 no His construct, lysed protein was purified by cation exchange using a 1 mL HiTrap SP HP (GE Healthcare) in protein buffer (50 mM Tris-HCl pH 7.0, 2 mM MgCl_2_, 1 mM EGTA and 10%) and eluted with a salt gradient from 50mM to 400mM NaCl. Protein was further purified using Streptactin affinity chromatography as per the other constructs. Purified CKAP2 was always used fresh for future experiments. Protein concentration was determined by absorbance at 288 or 506 nm with a DS-11 FX spectrophotometer (DeNovix, Inc.).

Tubulin was purified from bovine brains as previously described (Ashford and Hyman, 2006) with the modification of using Fractogel EMD SO3- (M) resin (Millipore-Sigma) instead of phosphocellulose. Tubulin was labeled using Atto-633 NHS-Ester (ATTO-TEC) and tetramethylrhodamine (TAMRA, Invitrogen) as described (Hyman et al., 1991). An additional cycle of polymerization/depolymerization was performed before use. Protein concentrations were determined using a DS-11 FX spectrophotometer (DeNovix, Inc.).

### *In vitro* Microscopy

The microscope set-up uses a Zeiss Axiovert Z1 microscope chassis and a 100x 1.45 NA Plan-apochromat objective lens. Total internal reflection fluorescence (TIRF) was achieved by coupling 488/561/637nm lasers to an iLas2 targeted laser illumination system (BioVision, Inc.) equipped with 360° TIRF. The objective was heated to 35°C with a CU-501 Chamlide lens warmer (Live Cell Instrument). Data was also acquired with a customized Zeiss Axio Observer 7 equipped with a Laser TIRF III and 405/488/561/638 nm lasers, and Alpha Plan-Apo 100x/1.46Oil DIC M27 Objective Heater 25.5/33 S1. Images on both systems were recorded on a Prime 95B CMOS camera (Photometrics) with a pixel size of 107 nm.

### Dynamic Microtubule Growth Assay

To visualize dynamic microtubules, we reconstituted microtubule growth off of GMPCPP double stabilized microtubule ‘seeds’(Gell et al., 2010). In short, cover glass was cleaned in acetone, sonicated in 50% methanol, sonicated in 0.5 M KOH, exposed to air plasma (Plasma Etch) for 3 min, then silanized by soaking in 0.2% Dichlorodimethylsilane (DDS) in n-Heptane. 5 μL flow channels were constructed using two pieces of silanized cover glasses (22 × 22 mm and 18 × 18 mm) held together with double-sided tape and mounted into custom-machined cover slip holders. GMPCPP seeds were prepared by polymerizing a 1:4 molar ratio of TAMRA labelled:unlabelled tubulin in the presence of guanosine-5’-[(*α*, *β*)-methyleno]triphosphate (GMPCPP, Jena Biosciences) in two cycles, as described by Gell et al. 2010. Channels were first incubated with anti-TAMRA antibodies (Invitrogen) and then blocked with 5% Pluronic F-127. Flow channels were washed 3x with BRB80 before incubating with GMPCPP seeds. On each day of experiments tubes of unlabelled and Atto-633 labelled tubulin was thawed and mixed at a 1:17 molar ratio and then sub-aliquoted and refrozen in liquid nitrogen. For consistency in microtubule growth dynamics, one sub-aliquot of tubulin was used for each experiment. Microtubule growth from GMPCPP seeds was achieved by incubating flow channels with tubulin in imaging buffer: BRB80, 1 mM GTP, 0.1 mg/mL BSA, 10 mM dithiothreitol, 250 nM glucose oxidase, 64 nM catalase, and 40 mM D-glucose. For low concertation tubulin experiments (50-500 nM), low retention tips and tubes were used.

### Turbidity Bulk-Phase Microtubule Polymerization Assay

The formation of microtubule polymer was followed as the absorbance at 350 nm using a Cary 300 Bio UV-Visible spectrophotometer or Cary 3500 UV-Vis spectrophotometer (Varian Inc., Agilent Technologies, Santa Clara, CA). Tubulin was polymerized at 36 °C or 4 °C in the presence or absence of CKAP2 in BRB80 containing 1 mM GTP and 1 mM MgCl_2_.

### Microtubule Nucleation Assay

Flow channels were constructed as previously described but anti-TAMRA antibody was replaced with Anti-ß3 tubulin antibody (BioLegend Poly18020 anti-Tubulin Beta 3). Microtubule nucleation was achieved by introducing 5-50 μM tubulin (1:17 molar ratio of Atto-633 labelled:unlabelled) in imaging buffer: into the flow channel. For CKAP2 positive experiments, freshly purified 0.2 μM CKAP2-mNG was introduced with 0.2-0.5 μM tubulin (1:17 molar ratio of Atto-633 labelled:unlabelled) in imaging buffer into flow channels. Tubulin was incubated 2 min, after which the total number of microtubules nucleated within one field of view (110 × 110 μm) was counted.

### Microtubule Depolymerization Assay

Assay channels were constructed as described above and incubated with 25% labelled TAMRA GMPCPP seeds. 0 μM or 1 μM CKAP2-mNG in imaging buffer without GTP was introduced into the channel. Channels were capped with nail polish to allow for longer imaging times. Microtubule seeds were imaged every 1 min during depolymerization for up to 10 hours.

### Preparation of Paclitaxel-Stabilized GDP Microtubules

A polymerization mixture was prepared with BRB80 + 32 μM tubulin + 1 mM GTP + 4 mM MgCl_2_ + 5% DMSO. The mixture was incubated on ice for 5 min, followed by incubation at 37°C for 30 min. The polymerized microtubules were diluted into prewarmed BRB80 + 10 μM paclitaxel, centrifuged at 110,000 rpm (199,000 × *g*) in an Airfuge (Beckman-Coulter), and resuspended in BRB80 + 10 μM paclitaxel (99.5+%, GoldBio)

### Curvature Recognition

TAMRA labelled paclitaxel microtubules were introduced into flow channels (Bechstedt et al., 2014). 5 nM CKAP2-mNG in imaging buffer with 10 μM paclitaxel without GTP was introduced into the channel. Quantification of the microtubule curvature, κ, and CKAP2-mNG signal on microtubules was analyzed using Kappa (Mary and Brouhard, 2019).

### Image Analysis and Software

For all *in vitro* data, image acquisition was controlled using MetaMorph (Molecular Devices) or ZEN 2.3 (Zeiss). Images were acquired from 2 sec to 1 min intervals.

All images were processed and analyzed using Fiji (Schindelin et al., 2012) (ImageJ). If needed, prior to analysis images were corrected for stage drift using a drift correct script (Hadim). Microtubule dynamics were analyzed using kymographs (for cell data and *in vitro* data). Growth and shrinkage rates were measured by manually drawing lines on kymographs and measuring the slope of growth or shrinkage.

Probability to nucleate was calculated by counting the number of seeds to nucleate a microtubule within 1 min over the total number of seeds within a field of view. Catastrophe frequency was measured by counting the total number of catastrophe events over the total time of all microtubule growth within a channel.

All functions were fitted and graphed with OriginPro2020 (OriginLab) or Python 3 (available at python.org) using a JupyterLab Notebook. Mean and standard deviation were calculated using Excel. Statistical analysis was performed using OriginPro2020. Images were linearly adjusted in brightness and contrast using Photoshop (Adobe). All figures were assembled using Illustrator (Adobe).

## Acknowledgements

We thank Gary Brouhard as well as members of the Brouhard and Bechstedt labs for critical reading of the manuscript.

## Author contributions

S.B. and T.M. designed the study, performed and analyzed experiments and wrote the manuscript.

## Data and materials availability

All data is available in the main text or the supplementary materials. Raw data and reagents will be made available upon request.

---

**Supplemental Figure 1.**
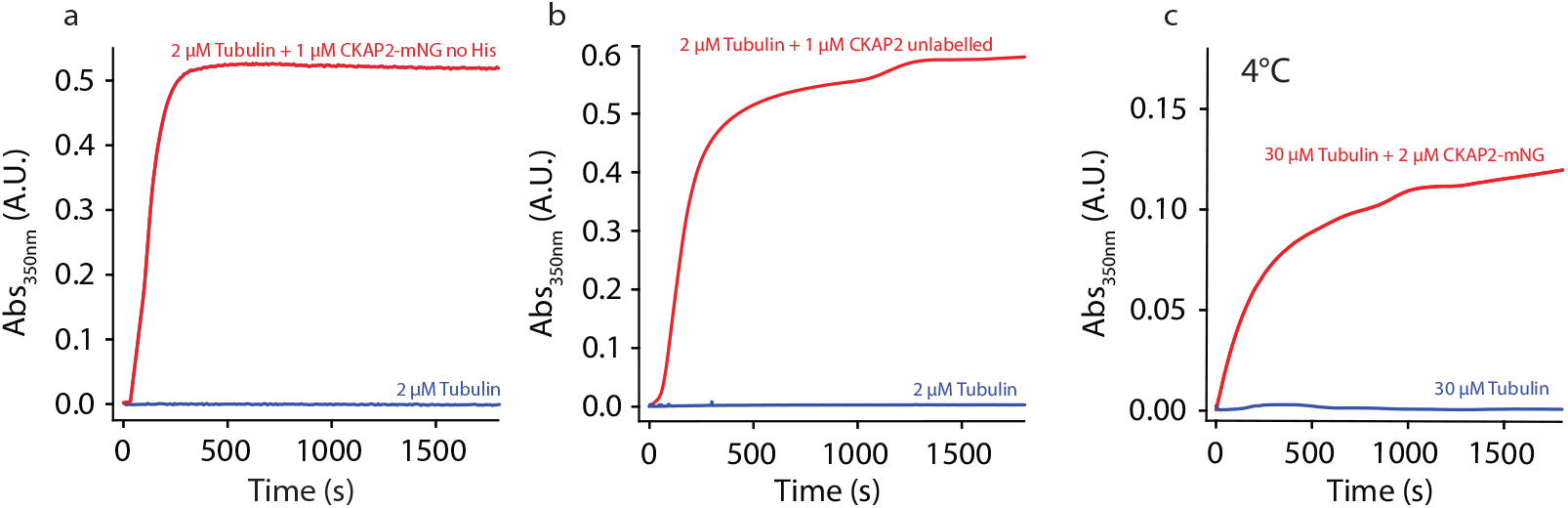
CKAP2 is an intrinsically disordered protein that increases microtubule assembly. (a) Turbidity assay with CKAP2-mNG no-His construct. (b) Turbidity assay with CKAP2 unlabeled (no-mNG). (c) Turbidity assay performed with CKAP2-mNG at 4°C.

**Supplemental Figure 2.**
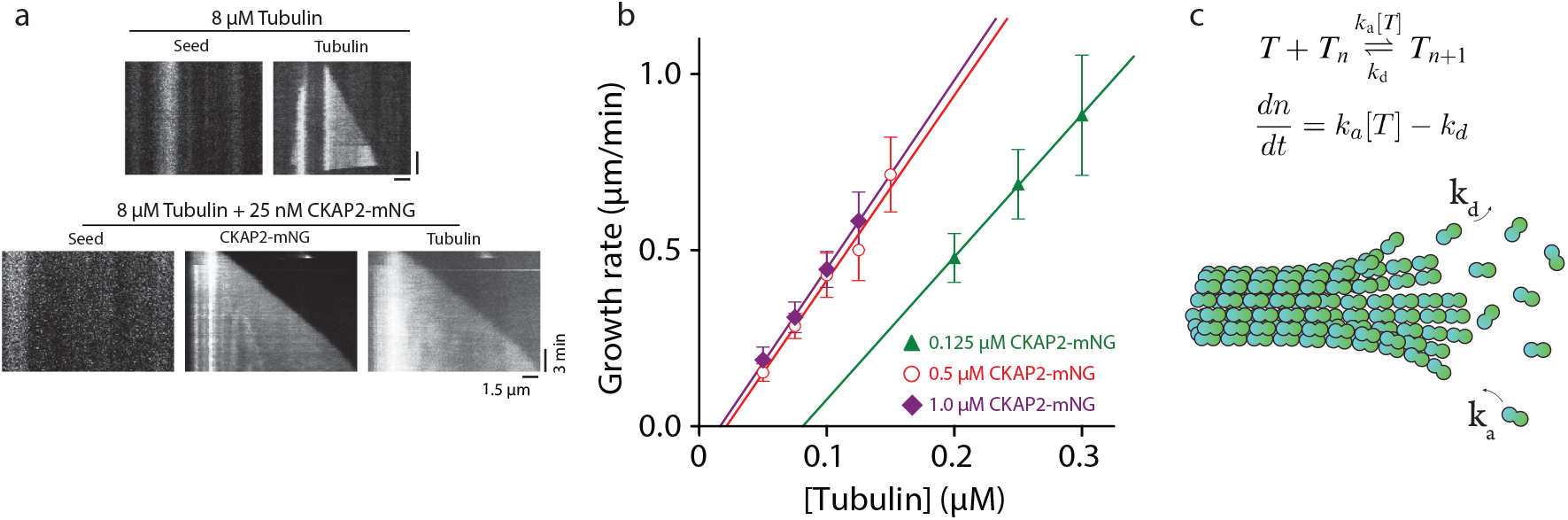
CKAP2 lowers the critical concentration of microtubule growth and speeds up microtubule assembly rates. **(a)** Representative kymograph of microtubule growth in the presence and absence of CKAP2-mNG. **(b)** Enlargement at low tubulin concentrations of microtubule growth rate plot from Figure 2C. **(c)** Schematic of linear polymer growth equation used to derive apparent on rate constant *k_a_*, the apparent off rate constant *k_d_*, and critical concentration *C_c_* (Oosawa et al., 1975). *k_a_* was calculated from the slope, *k_d_*, the y-intercept, and *C_c_* the x-axis of the linear regression fit.

**Supplemental Figure 3.**
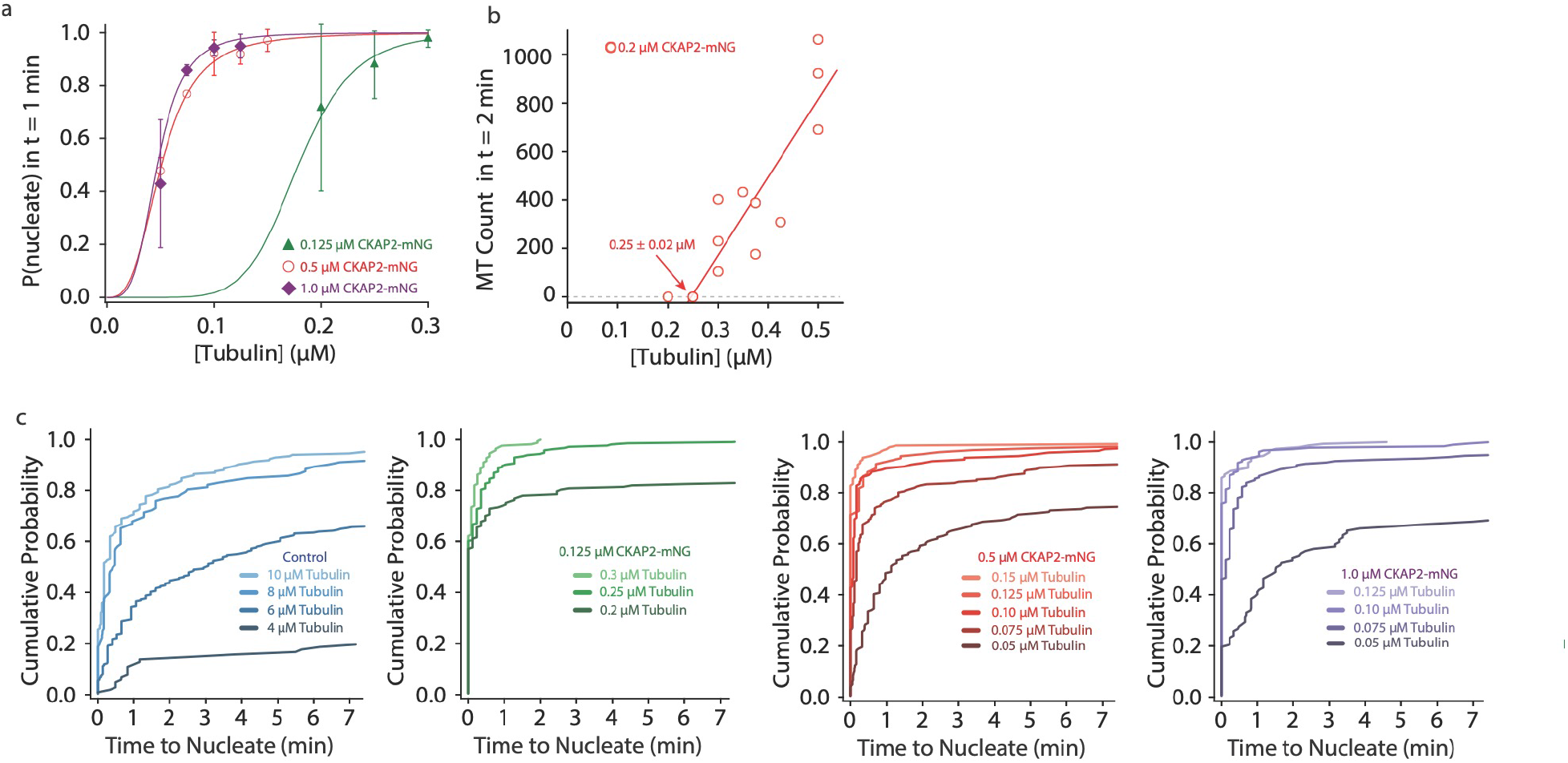
CKAP2 increases templated and spontaneous microtubule nucleation. **(a)** Enlargement at low tubulin concentrations of probability to nucleate plot from Figure 3C. **(b)** Enlargement at low tubulin concentrations of spontaneous nucleation in the presence of 0.2 μM CKAP2 from Figure 3E. **(c)** Plots of the cumulative probability distributions for time to nucleate as a function of tubulin concentration, same *n* as Fig 3C.

**Supplemental Figure 4.**
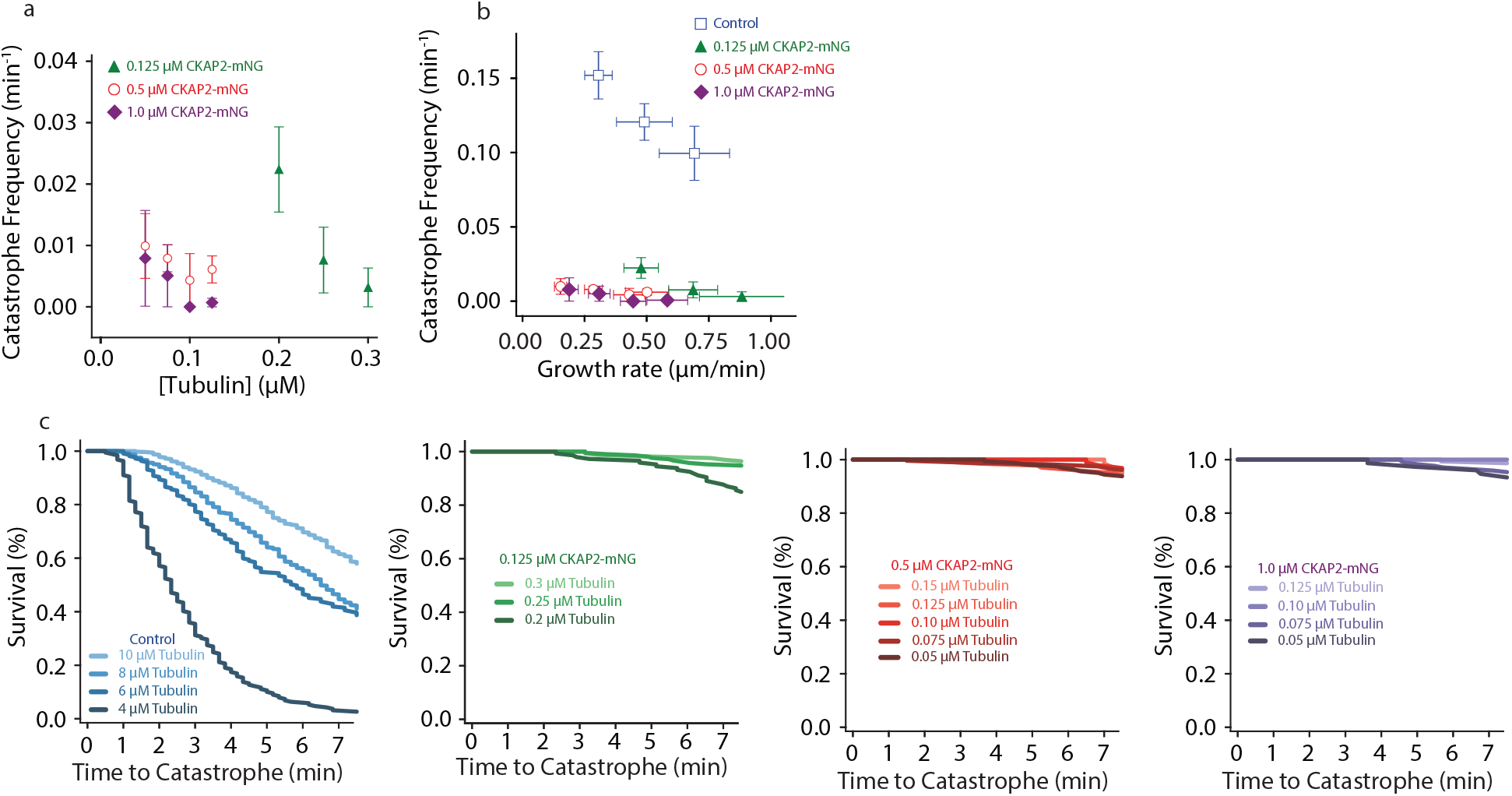
CKAP2 suppresses catastrophe. **(a)** Enlargement at low tubulin concentrations of catastrophe frequency plot Figure 4B. **(b)** Plot of tubulin growth rate versus catastrophe frequency. **(c)** Plot of the cumulative microtubule survival % as a function of time, same *n* as Figure 4B.

**Supplemental Figure 5.**
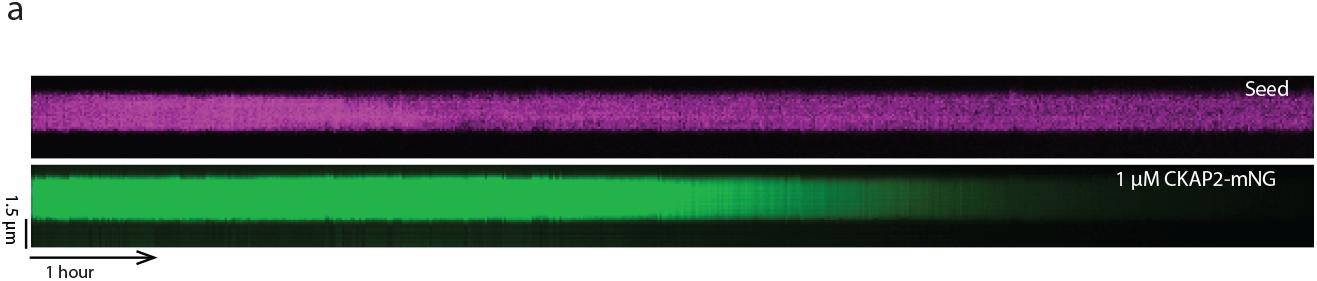
CKAP2 recruits tubulin, recognizes lattice curvature, but does not catalyze microtubule depolymerization. **(a)** GMPCPP ‘seed’ depolymerization kymograph for 1 μM CKAP2-mNG over 10 hours

## References

Aguirre-Portolés, C., Bird, A.W., Hyman, A., Cañamero, M., Pérez de Castro, I., and Malumbres, M. (2012). Tpx2 controls spindle integrity, genome stability, and tumor development. Cancer Res 72, 1518–1528.

Aher, A., Kok, M., Sharma, A., Rai, A., Olieric, N., Rodriguez-Garcia, R., Katrukha, E.A., Weinert, T., Olieric, V., Kapitein, L.C., et al. (2018). CLASP Suppresses Microtubule Catastrophes through a Single TOG Domain. Developmental Cell.

Ashford, A.J., and Hyman, A.A. (2006). Chapter 22 - Preparation of Tubulin from Porcine Brain. In Cell Biology (Third Edition), J.E. Celis, ed. (Burlington: Academic Press), pp. 155–160.

Ayaz, P., Munyoki, S., Geyer, E.A., Piedra, F.-A., Vu, E.S., Bromberg, R., Otwinowski, Z., Grishin, N.V., Brautigam, C.A., and Rice, L.M. (2014). A tethered delivery mechanism explains the catalytic action of a microtubule polymerase. ELife Sciences 3, e03069.

Bechstedt, S., and Brouhard, G.J. (2012). Doublecortin recognizes the 13-protofilament microtubule cooperatively and tracks microtubule ends. Dev. Cell 23, 181–192.

Bechstedt, S., Lu, K., and Brouhard, G.J. (2014). Doublecortin Recognizes the Longitudinal Curvature of the Microtubule End and Lattice. Curr. Biol.

Brouhard, G.J., Stear, J.H., Noetzel, T.L., Al-Bassam, J., Kinoshita, K., Harrison, S.C., Howard, J., and Hyman, A.A. (2008). XMAP215 is a processive microtubule polymerase. Cell 132, 79–88.

Case, C.M., Sackett, D.L., Wangsa, D., Karpova, T., McNally, J.G., Ried, T., and Camps, J. (2013). CKAP2 Ensures Chromosomal Stability by Maintaining the Integrity of Microtubule Nucleation Sites. PLoS ONE 8, e64575.

Consolati, T., Locke, J., Roostalu, J., Chen, Z.A., Gannon, J., Asthana, J., Lim, W.M., Martino, F., Cvetkovic, M.A., Rappsilber, J., et al. (2020). Microtubule Nucleation Properties of Single Human γTuRCs Explained by Their Cryo-EM Structure. Developmental Cell 53, 603–617.e8.

Desai, A., and Mitchison, T.J. (1997). MICROTUBULE POLYMERIZATION DYNAMICS. Annual Review of Cell and Developmental Biology 13, 83–117.

Dosztányi, Z., Csizmók, V., Tompa, P., and Simon, I. (2005). The pairwise energy content estimated from amino acid composition discriminates between folded and intrinsically unstructured proteins. J. Mol. Biol. 347, 827–839.

Ertych, N., Stolz, A., Stenzinger, A., Weichert, W., Kaulfuß, S., Burfeind, P., Aigner, A., Wordeman, L., and Bastians, H. (2014). Increased microtubule assembly rates influence chromosomal instability in colorectal cancer cells. Nat Cell Biol 16, 779–791.

Gard, D.L., and Kirschner, M.W. (1987). A microtubule-associated protein from Xenopus eggs that specifically promotes assembly at the plus-end. J Cell Biol 105, 2203–2215.

Gell, C., Bormuth, V., Brouhard, G.J., Cohen, D.N., Diez, S., Friel, C.T., Helenius, J., Nitzsche, B., Petzold, H., Ribbe, J., et al. (2010). Microtubule dynamics reconstituted in vitro and imaged by single-molecule fluorescence microscopy. Methods Cell Biol 95, 221–245.

Gergely, F., Draviam, V.M., and Raff, J.W. (2003). The ch-TOG/XMAP215 protein is essential for spindle pole organization in human somatic cells. Genes Dev 17, 336–341.

Guo, Q., Song, Y., Hua, K., and Gao, S. (2017). Involvement of FAK-ERK2 signaling pathway in CKAP2-induced proliferation and motility in cervical carcinoma cell lines. Scientific Reports 7, 2117.

Hayashi, T., Ohtsuka, M., Okamura, D., Seki, N., Kimura, F., Shimizu, H., Yoshidome, H., Kato, A., Yoshitomi, H., Furukawa, K., et al. (2014). Cytoskeleton-associated protein 2 is a potential predictive marker for risk of early and extensive recurrence of hepatocellular carcinoma after operative resection. Surgery 155, 114–123.

Hong, K.U., Park, Y.S., Seong, Y.-S., Kang, D., Bae, C.-D., and Park, J. (2007). Functional Importance of the Anaphase-Promoting Complex-Cdh1-Mediated Degradation of TMAP/CKAP2 in Regulation of Spindle Function and Cytokinesis. Mol. Cell. Biol. 27, 3667–3681.

Hong, K.U., Choi, Y.-B., Lee, J.-H., Kim, H.-J., Kwon, H.-R., Seong, Y.-S., Kim, H.T., Park, J., Bae, C.-D., and Hong, K.-M. (2008). Transient phosphorylation of tumor associated microtubule associated protein (TMAP)/cytoskeleton associated protein 2 (CKAP2) at Thr-596 during early phases of mitosis. Exp Mol Med 40, 377–386.

Hong, K.U., Kim, H.-J., Bae, C.-D., and Park, J. (2009). Characterization of mitosis-specific phosphorylation of tumor-associated microtubule-associated protein. Exp. Mol. Med. 41, 832–840.

Howard, J., and Hyman, A.A. (2007). Microtubule polymerases and depolymerases. Curr Opin Cell Biol 19, 31–35.

Hyman, A., Drechsel, D., Kellogg, D., Salser, S., Sawin, K., Steffen, P., Wordeman, L., and Mitchison, T. (1991). Preparation of modified tubulins. Methods Enzymol 196, 478–485.

Hyman, A.A., Salser, S., Drechsel, D.N., Unwin, N., and Mitchison, T.J. (1992). Role of GTP hydrolysis in microtubule dynamics: information from a slowly hydrolyzable analogue, GMPCPP. Mol Biol Cell 3, 1155–1167.

Jin, Y., Murakumo, Y., Ueno, K., Hashimoto, M., Watanabe, T., Shimoyama, Y., Ichihara, M., and Takahashi, M. (2004). Identification of a mouse cytoskeleton-associated protein, CKAP2, with microtubule-stabilizing properties. Cancer Science 95, 815–821.

Kapoor, T.M. (2017). Metaphase Spindle Assembly. Biology 6, 8.

Kollman, J.M., Polka, J.K., Zelter, A., Davis, T.N., and Agard, D.A. (2010). Microtubule nucleating gamma-TuSC assembles structures with 13-fold microtubule-like symmetry. Nature 466, 879–882.

Kovar, D.R., Harris, E.S., Mahaffy, R., Higgs, H.N., and Pollard, T.D. (2006). Control of the Assembly of ATP- and ADP-Actin by Formins and Profilin. Cell 124, 423–435.

Lawrence, E.J., Arpag, G., Norris, S.R., and Zanic, M. (2018). Human CLASP2 specifically regulates microtubule catastrophe and rescue. MBoC 29, 1168–1177.

Majumdar, S., Kim, T., Chen, Z., Munyoki, S., Tso, S.-C., Brautigam, C.A., and Rice, L.M. (2018). An isolated CLASP TOG domain suppresses microtubule catastrophe and promotes rescue. MBoC 29, 1359–1375.

Mary, H., and Brouhard, G.J. (2019). Kappa (κ): Analysis of Curvature in Biological Image Data using B-splines. BioRxiv 852772.

Mitchison, T., and Kirschner, M. (1984). Dynamic instability of microtubule growth. Nature 312, 237–242.

Moores, C.A., Perderiset, M., Kappeler, C., Kain, S., Drummond, D., Perkins, S.J., Chelly, J., Cross, R., Houdusse, A., and Francis, F. (2006). Distinct roles of doublecortin modulating the microtubule cytoskeleton. EMBO J 25, 4448–4457.

Moritz, M., Braunfeld, M.B., Sedat, J.W., Alberts, B., and Agard, D.A. (1995). Microtubule nucleation by gamma tubulin-containing rings in the centrosome. Nature 378, 638–640.

Norholm, M.H. (2010). A mutant Pfu DNA polymerase designed for advanced uracil-excision DNA engineering. BMC Biotechnol 10, 21.

Odde, D.J. (1997). Estimation of the diffusion-limited rate of microtubule assembly. Biophys J 73, 88–96.

Oosawa, F., Asakura, S., Asakura, S., and 1927- (1975). Thermodynamics of the polymerization of protein (New York and London: Academic Press).

Petry, S., and Vale, R.D. (2015). Microtubule nucleation at the centrosome and beyond. Nat Cell Biol 17, 1089–1093.

Petry, S., Pugieux, C., Nédélec, F.J., and Vale, R.D. (2011). Augmin promotes meiotic spindle formation and bipolarity in Xenopus egg extracts. PNAS 108, 14473–14478.

Piehl, M., Tulu, U.S., Wadsworth, P., and Cassimeris, L. (2004). Centrosome maturation: Measurement of microtubule nucleation throughout the cell cycle by using GFP-tagged EB1. PNAS 101, 1584–1588.

Reid, T.A., Schuster, B.M., Mann, B.J., Balchand, S.K., Plooster, M., McClellan, M., Coombes, C.E., Wadsworth, P., and Gardner, M.K. (2016). Suppression of microtubule assembly kinetics by the mitotic protein TPX2. J Cell Sci 129, 1319–1328.

Roostalu, J., and Surrey, T. (2017). Microtubule nucleation: beyond the template. Nature Reviews Molecular Cell Biology 18, 702–710.

Roostalu, J., Cade, N.I., and Surrey, T. (2015). Complementary activities of TPX2 and chTOG constitute an efficient importin-regulated microtubule nucleation module. Nature Cell Biology 17, 1422–1434.

Saxton, W.M., Stemple, D.L., Leslie, R.J., Salmon, E.D., Zavortink, M., and McIntosh, J.R. (1984). Tubulin dynamics in cultured mammalian cells. J Cell Biol 99, 2175–2186.

Schindelin, J., Arganda-Carreras, I., Frise, E., Kaynig, V., Longair, M., Pietzsch, T., Preibisch, S., Rueden, C., Saalfeld, S., Schmid, B., et al. (2012). Fiji: an open-source platform for biological-image analysis. Nat Methods 9, 676–682.

Seki, A., and Fang, G. (2007). CKAP2 Is a Spindle-associated Protein Degraded by APC/C-Cdh1 during Mitotic Exit. J. Biol. Chem. 282, 15103–15113.

Strome, S., Powers, J., Dunn, M., Reese, K., Malone, C.J., White, J., Seydoux, G., and Saxton, W. (2001). Spindle Dynamics and the Role of γ-Tubulin in Early Caenorhabditis elegans Embryos. Mol Biol Cell 12, 1751–1764.

Thawani, A., Kadzik, R.S., and Petry, S. (2018). XMAP215 is a microtubule nucleation factor that functions synergistically with the γ-tubulin ring complex. Nat Cell Biol 20, 575–585.

Tsuchihara, K., Lapin, V., Bakal, C., Okada, H., Brown, L., Hirota-Tsuchihara, M., Zaugg, K., Ho, A., Itie-YouTen, A., Harris-Brandts, M., et al. (2005). Ckap2 Regulates Aneuploidy, Cell Cycling, and Cell Death in a p53-Dependent Manner. Cancer Res 65, 6685–6691.

Uehara, R., Nozawa, R., Tomioka, A., Petry, S., Vale, R.D., Obuse, C., and Goshima, G. (2009). The augmin complex plays a critical role in spindle microtubule generation for mitotic progression and cytokinesis in human cells. PNAS 106, 6998–7003.

Uversky, V.N., and Dunker, A.K. (2013). The case for intrinsically disordered proteins playing contributory roles in molecular recognition without a stable 3D structure. F1000 Biol Rep 5.

Wieczorek, M., Bechstedt, S., Chaaban, S., and Brouhard, G.J. (2015). Microtubule-associated proteins control the kinetics of microtubule nucleation. Nat Cell Biol 17, 907–916.

Wühr, M., Freeman, R.M., Jr, Presler, M., Horb, M.E., Peshkin, L., Gygi, S., and Kirschner, M.W. (2014). Deep Proteomics of the Xenopus laevis Egg using an mRNA-derived Reference Database. Current Biology : CB 24, 1467.

Yu, G., Lee, Y.-C., Cheng, C.-J., Wu, C.-F., Song, J.H., Gallick, G.E., Yu-Lee, L.-Y., Kuang, J., and Lin, S.-H. (2015). RSK Promotes Prostate Cancer Progression in Bone through ING3, CKAP2, and PTK6-Mediated Cell Survival. Mol Cancer Res 13, 348–357.

Yu, N., Signorile, L., Basu, S., Ottema, S., Lebbink, J.H.G., Leslie, K., Smal, I., Dekkers, D., Demmers, J., and Galjart, N. (2016). Isolation of Functional Tubulin Dimers and of Tubulin-Associated Proteins from Mammalian Cells. Current Biology 26, 1728–1736.

Zanic, M., Widlund, P.O., Hyman, A.A., and Howard, J. (2013). Synergy between XMAP215 and EB1 increases microtubule growth rates to physiological levels. Nature Cell Biology 15, 688–693.

